# Defining mutational signatures of lung cancer-associated carcinogens through *in vitro* exposure of human airway epithelial cells

**DOI:** 10.64898/2026.03.05.707509

**Authors:** Natasha Q. Gurevich, Darren J. Chiu, Masanao Yajima, Jonathan Huggins, Sarah A. Mazzilli, Joshua D. Campbell

**Affiliations:** Section of Computational Biomedicine, Department of Medicine, Boston University Chobanian & Avedisian School of Medicine, Boston, Massachusetts; Bioinformatics Program, Boston University, Boston, Massachusetts; Department of Mathematics & Statistics, Boston University, Boston, Massachusetts; Faculty of Computing and Data Sciences, Boston University, Boston, Massachusetts

**Author notes:** **Corresponding authors:** Joshua D. Campbell, 72 E Concord St Boston, MA 02118, Sarah A. Mazzilli, 72 E Concord St Boston, MA 02118. M. Yajima in now an employee at Takeda Pharmaceuticals. These authors serve as co-senior authors (Sarah A. Mazzilli, Joshua D. Campbell).

**Keywords:** Mutational signatures, cancer genomics, lung cancer, bioinformatics

## Abstract

While distinct environmental exposures imprint unique mutational signatures on cancer genomes, the specific causal patterns for many known carcinogens remain uncharacterized in relevant human tissues. To address this gap, we developed a novel, physiologically relevant system that uses a combination of airway epithelial cells and whole genome sequencing to characterize mutational patterns induced by genotoxic carcinogens associated with lung cancer. After validating the platform’s accuracy by successfully recapturing the known signature for Benzo(a)pyrene (BaP), we used this system to gain detailed insights into the types of mutations that occur with exposure to N-nitrosotris-(2-chloroethyl) urea (NTCU) and 4-(methylnitrosamino)-1-(3-pyridyl)-1-butanone (NNK), genotoxic compounds that induce lung squamous cell carcinoma and lung adenocarcinoma in mouse models, respectively. Cells exposed to NTCU had significantly more somatic SNVs compared to control samples. An average of 82.3% of mutations in NTCU samples were attributed to a novel mutational signature distinct from those in the COSMIC database but highly correlated with recent *in vivo* mouse models. In contrast, NNK exposure did not demonstrate a distinct mutational pattern above background at both high and low concentrations. Ultimately, this *in vitro* system provides a robust platform to define causal links between environmental exposures and mutational patterns in lung cancer mutagenesis.

**Statement of Significance:** *In vitro* exposure of N-nitrosotris-(2-chloroethyl) urea to airway epithelial cells revealed a distinct mutational signature.

## INTRODUCTION

Exogenous exposures and endogenous biological processes can induce mutations in the genome. Distinct exposures leave unique, probabilistic footprints of DNA damage, termed mutational signatures [1,2]. These mutations may include single or double base substitutions (SBS, DBS), insertions or deletions (INDELs), genome rearrangements and structural variants (SVs), or copy number variants (CNVs). Because a single tumor typically accumulates mutations from multiple active processes, its overall mutational catalogue serves as a historical record of environmental exposures and provides critical insight into cancer etiology [1–4].

To date, over 100 mutational signatures have been characterized in the Catalogue of Somatic Mutations in Cancer (COSMIC) database [5]. While specific etiologies—such as cigarette smoke, UV-light, and aristolochic acid—have been identified for many signatures, numerous others remain uncharacterized. While clinical history can be examined to gain insights into possible chemical exposures, more direct methods to characterize the mutational signatures associated with specific chemical exposures are necessary to accurately and rapidly identify putative etiologies. High-throughput DNA sequencing such as whole-genome sequencing (WGS) can comprehensively characterize the set of genomic alterations within a sample in an unbiased fashion at single-base resolution. While this technology has been widely adopted to study human tumors, more recent studies have begun to use WGS of model systems in the field of toxicology to characterize mutational patterns in response to carcinogen exposure [6–8].

N-nitrosotris-(2-chloroethyl) urea (NTCU) and 4-(methylnitrosamino)-1-(3-pyridyl)-1-butanone (NNK) are two known carcinogens associated with the development of lung cancer that do not have known mutational signatures. Specifically, NTCU has been shown to induce premalignant lesions that develop into lung squamous cell carcinoma (SCC) in SWR/J mice [9].

RNA sequencing has revealed a high degree of similarity in driver mutations and expression profiles between human lung SCC and mouse lung SCC induced by NTCU treatment [10] and the histological progression of airway premalignant lesions to invasive SCC in humans has been replicated in these mouse models [11]. However, the exact mechanisms by which NTCU induces SCC in mice remains unknown and the mutational footprint of NTCU has not been adequately defined. NNK is a potent carcinogen found in tobacco smoke and can be used to induce lung adenoma and adenocarcinoma in mice [12–14]. Although this carcinogen induces epigenetic alterations and affects various cancer associated signaling pathways, it has no documented mutational signature [15,16]. Benzo(a)pyrene (BaP) is a polycyclic aromatic hydrocarbon formed during incomplete combustion of organic material. It is found in tobacco smoke and has a mutational phenotype well defined to represent smoking [1,4,6].

We have developed a novel physiologically relevant system that uses a combination of airway epithelial cells and WGS to characterize mutational patterns induced by genotoxic carcinogens associated with lung cancer. Cells were treated with NTCU, NNK, and BaP carcinogens *in vitro*, potent lung carcinogens widely utilized in mouse models of squamous cell carcinoma and adenocarcinoma. We found distinct patterns of mutations associated with NTCU treatment, including a higher number of mutations compared to controls and a mutational signature highly active in only NTCU samples that has not been documented in the COSMIC database. In contrast, NNK-treated samples did not have a distinct mutational signature suggesting it has limited ability to induce specific types of mutations in this experimental context.

## MATERIALS AND METHODS

### Cell Culture and Carcinogen Exposure

Human immortalized (SV40 and hTERT) tracheobronchial epithelial cells (AALEs) were maintained in human bronchial/tracheal epithelial cell media (Cell Application, Cat# 511-500). Cells were treated with two solvent controls: 0.1% DMSO and 1% acetone, and three carcinogens: benzo(a)pyrene (BaP; Sigma, Cat# B1760), N-nitrosotris-(2-chloroethyl)urea (NTCU; Toronto Research Chemicals), and 4-(methylnitrosamino)-1-(3-pyridyl)-1-butanone (NNK; Toronto Research Chemicals, Cat# M325750). Exposures were performed at the indicated concentrations in the presence or absence of 10ug/ml liver microsomes (Sigma, Cat# M9066) and 0.1mM NADPH (Sigma, Cat# 10107824001) for metabolic activation.

### Viability and DNA Damage Assessment

Cell viability was assessed using the CellTiter-Glo® Luminescent Cell Viability Assay (Promega) according to the manufacturer’s instructions. DNA damage was quantified via long amplicon quantitative PCR (LA-QPCR), as previously described [17]. Briefly, total genomic DNA was extracted using the QIAamp DNA Kit (Qiagen, Cat# 51304). A total of 10 ng DNA was used as a template to amplify long and short amplicons of the mitochondrial genomes using Herculase II Fusion DNA Polymerase (Agilent, Cat# 600675). Primer sequences are provided in **Supplementary Table 1**. Amplified DNA was quantified using the Quant-iT™ PicoGreen™ dsDNA Assay Kit (Fisher Scientific, Cat# P7589). DNA lesion frequency was calculated by comparing the relative amplification of treated samples to that of untreated controls.

### Generation of Carcinogen-Exposed cell lines

AALE cells were cultured in 6-well plates and treated with carcinogens *in vitro* for seven days. The treatment conditions were determined by cytotoxicity and the DNA damage response via the LA-QPCR assay. These cells have expressions of members of the cytochrome p450 family necessary for processing carcinogens in cigarette smoke such as CYP1A1 and CYP1B1 and thus are generally competent to bioactivate carcinogens. After 7 days, cells were re-plated and exposed to carcinogens. Control samples consisted of AALE cells treated with media control (N =3), 0.1% DMSO (N = 3), and 1% acetone (N = 3). The carcinogen-exposed samples consisted of AALE cells treated with 5 μM BaP in 0.1% DMSO supplemented with 10 ug/ml liver microsome and 0.1 mM NADPH for activation (N = 4); 0.5 μM NTCU in 1% acetone (N = 4); 0.01 μM NNK in 0.1% DMSO (N = 4); and 1.0 μM NNK in 0.1% DMSO (N = 4). This process repeated for four cycles, totaling four weeks of exposure. Following the final exposure, cells were harvested and stained with Calcein Blue (Thermo Fisher Scientific, Cat# C1429) for a viability-based flow cytometric sorting. Live single cells were sorted into 96-well plates using a Beckman Coulter MoFlo Astrios sorter. Individual clones were expanded and transferred to 6-well plates upon reaching approximately 80% confluence for further expansions.

### Sequencing and alignment

DNA was isolated using the QIAamp DNA kit (Qiagen, Cat# 51304) from replicates from each condition, a total of 25 samples, and sent to BGI Americas Corporation for WGS using the DNBseq platform with 100 base-pair paired-end reads and a minimum depth of 30x. The raw unmapped reads in FASTQ format were first converted to the uBAM format using Picard’s FastqToSam tool v2.25.2 [18]. GATK v4.2.6.1 was used for data preprocessing and variant discovery using the Best Practices workflow [19]. Briefly, BWA was used to map reads to the genome and GATK was used to mark duplicates and recalibrates base quality scores which provided analysis-ready reads in the BAM format.

### Somatic mutation calling

The GATK Mutect2 tool in paired tumor-normal mode was then used to call mutations for all control and treated samples [20]. For the samples treated with a carcinogen, a control sample with the same solvent as the treatment sample was used for the “matched normal”. To call mutations in an untreated control sample, another control sample with no solvent was used for the matched normal. A panel of normals was created using the CreateSomaticPanelOfNormals tool from GATK and included in each run. Variant calling for the treatment samples was performed with a panel of normals consisting of all nine control samples. Variant calling for each control sample was performed using a unique panel of normals that contained all eight other control samples. The musicatk package [21] was then used to extract variants from the VCF files and build standard mutation count tables for single base substitutions (SBS), double base substitutions (DBS), and indels (INDEL).

### Variant calling for subclonal analysis

To identify the germline single nucleotide polymorphisms (SNPs) and clonal somatic variants present in the parental line, the haplotypecaller-gvcf-gatk4 workflow was used to run the GATK4 HaplotypeCaller tool on individual samples according to GATK Best Practices [22]. Standard settings were used except GVCF-negative mode was used to directly call variants and output a VCF file for each sample. Variants present in only one sample were removed as these were likely somatic variants intrinsic to the single-cell clone. Similarly, variants present in all 9 samples were filtered out, as they were considered clonal mutations from the parental cell line or germline SNPs of the individual from which the cell line was derived and removed. The remaining variants present in 2 to 8 samples were used to perform PCA analysis and examine subclonal structure within the untreated control samples. The identification of these distinct subclonal groups informed our strategy to utilize a comprehensive panel or normals during somatic variant calling to effectively filter out this subclone-specific background.

### Single base substitution signature analysis

Single base substitutions were extracted from each sample to produce a mutation count table using the musicatk R package v2.0.0 [21]. The number of signatures that should be discovered in the dataset was determined using the “k value assist” tool in the musicatk package and validated with *de novo* NMF prediction with SigProfilerExtractor v1.1.16 [23]. We then used the NMF-based workflow in musicatk to discover three SBS signatures. The predicted signatures were compared to those from the COSMIC V3 database using a cosine similarity threshold of 0.8. The process detailed above for SBS signature analysis was repeated to explore double base substitution and indel signatures.

## RESULTS

### Establishing mutations using tobacco smoking related carcinogens in airway epithelial cells

To establish an *in vitro* system capable of capturing carcinogen-induced mutational signatures, cultured immortalized human tracheobronchial epithelial (AALE) cells were exposed to specific genotoxins at concentrations designed to balance measurable DNA damage with cellular viability (targeting 40-60% viability over seven days). We first optimized the metabolic activation requirements and dosing for each compound using long amplicon quantitative PCR (LA-QPCR) to assess mitochondrial DNA lesions. A concentration of 0.5 μM yielded the target viability for NTCU (**Figure 1A**). LA-QPCR indicated that NTCU alone causes more mitochondrial DNA damage than NTCU with liver microsome (**Figure 1B**). Conversely, BaP required metabolic activation, causing more damage in the presence of liver microsomes, leading us to select a 5 μM dose supplemented with microsomes (**Figure S1A-B**). For NNK, cytotoxicity analysis revealed minimal impact on viability, and LA-QPCR showed limited mitochondrial DNA damage regardless of microsome addition (**Figure S1C-D**). Consequently, cells were treated with NNK alone at two distinct concentrations: 0.01 μM and 1.0 μM.

**Figure 1.**
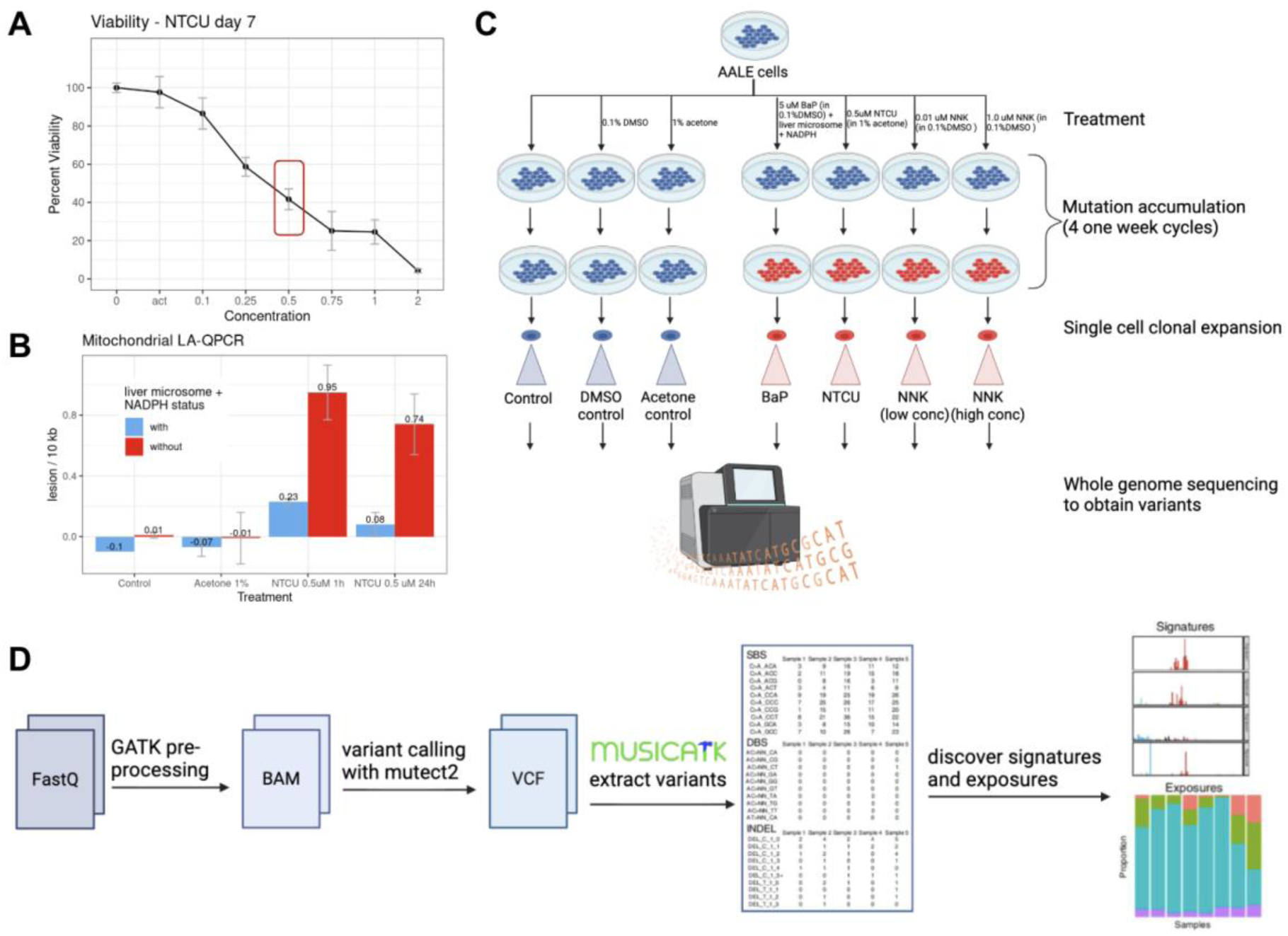
Experimental design for in vitro assessment of mutational signatures. **(A)** A cytotoxicity assay was used to determine the concentration of NTCU that results in 40-60% viability for seven days. A concentration of 0.5 μM was selected. **(B)** LA-QPCR was used to measure the DNA damage response with and without liver microsomes. NTCU alone causes more mitochondrial DNA damage than NTCU with liver microsome. **(C)** Human airway cells were cultured and treated in seven different conditions. Four of these conditions were treated with known genotoxic carcinogens from cigarette smoke or that can produce cancer in mouse models, including BaP, NTCU, high concentration NNK, and low concentration NNK. Three groups served as controls, including no solvent control, acetone solvent, and DMSO solvent. Mutations were allowed to accumulate for four one-week cycles. Single cells were sorted and individual clones were expanded and transferred to 6-well plates upon reaching approximately 80% confluence for further expansions. DNA was isolated and whole genome sequencing was performed. Four replicates for each of the four treatment groups and three replicates for each of the three control groups were sequenced. **(D)** Variant calling was performed using mutect2, following preprocessing of sequencing data using GATK best practices. Musicatk was used to extract variants from the resulting VCF files and then used to discover signatures and exposures.

Following this optimization, the carcinogen treatments and control exposures were repeated for four one-week cycles to allow for robust mutation accumulation before single-cell clonal expansion and whole-genome sequencing (**Figure 1C**). Control samples consisted of AALE cells with media control (N =3), AALE cells with 0.1% DMSO (control substrate for NNK/BaP) (N = 3), and AALE cells with 1% acetone (control substrate for NTCU) (N = 3). Exposure, or treated, samples consisted of AALE cells with 5 μM BaP in 0.1% DMSO, liver microsome, and NADPH for activation (N = 4); AALE cells with 0.5 μM NTCU in 1% acetone (N = 4); AALE cells with 0.01 μM NNK in 0.1% DMSO (N = 4); and AALE cells with 1.0 μM NNK in 0.1% DMSO (N = 4). WGS using the DNBseq platform with 100 base-pair paired-end reads was performed and the musicatk package was used for analysis following variant calling (**Methods**). We used the NMF-based workflow in musicatk to predict mutational signatures across all samples collectively. Single base substitution (SBS), double base substitution (DBS), and indel (INDEL) signatures were predicted (**Figure 1D**).

### Baseline mutational profiles in untreated AALE samples

An initial analysis of the single base substitutions in the control samples was performed to assess background mutation rates in this system without carcinogenic exposure. All nine controls have very similar patterns of mutation types (**Figure S2A**). The pattern is spread out over all six single base substitution types with some specific biases, such as T[C>A]T, T[C>T]T, and T[T>C]T. The average single nucleotide variant (SNV) mutation burden was 2,925.2. The higher mean was largely due to three samples with over four thousand mutations while the other control samples ranged between 1,706 and 2,223 (**Figure S2A**). Of the 28,892 unique variants in the control samples, 28,495 of them (98.6%) occur in only one sample while only 378 (1.3%) occurred in two or more samples, confirming clonal independence (**Figure S2B**). To explore potential parental subclone structure, HaplotypeCaller was used to find all variants in the control samples [22]. Variants observed in 2 to 8 of the 9 samples were analyzed with principal component analysis (PCA) to explore potential subclonal architecture (**Figure S2C**). Control samples fell into three broad groups based on the variation from PC1 and PC2. The first group was separated on PC1 and contained acetone controls 1 and 2, and DMSO controls 2 and 3. The second and third groups were separated along PC2 with group two containing standard controls 2 and 3 and group 3 containing standard control 1, DMSO control 1, and acetone control 3. These three samples are the samples that had a higher SNV mutation burden (**Figure S2A**). Germline SNPs and clonal somatic variants present in all control samples were also explored. Most variants (72.67%) were found in all nine controls, confirming the samples were derived from the same ancestral clone (**Figure S2D**). Because different control samples exhibited distinct subclonal architectures, utilizing a single matched control for variant calling risks confounding the analysis with private, subclone-specific mutations. We therefore implemented a comprehensive panel of normals to mitigate this risk by robustly filtering out background subclonal variants and isolating the true mutations induced by the carcinogen exposures.

### Distinct SBS patterns exist in AALEs treated with NTCU and BaP

We next explored the mutational profiles and mutational loads of the samples treated with BaP, NTCU, and NNK. BaP-, NTCU-, and NNK-exposed samples had distinct mutational profiles (**Figure 2A; Figure S3**). Mutations in BaP samples were dominated by C>A mutations, specifically at the C[C>A]C context. NTCU samples had mutations occurring in all six SBS categories with elevated levels of C>T and C>A mutations. NNK samples had the least specific pattern, with mutations occurring across all contexts in both the high and low dose treatment groups. Cosine correlation was used to quantify the similarities and differences between all samples (**Figure 2B**). The mutational profiles of samples within the same treatment group were all highly correlated across replicates (cosine > 0.9), demonstrating that each treatment produced a consistent pattern of mutations. For the NNK group, this includes consistency across both the high and low dose treatment. The mutational pattern of BaP and NTCU were distinct from the mutational patterns in the control samples (cosine < 0.85), while the pattern of NNK samples was highly correlated with that of the controls (cosine > 0.9).

**Figure 2.**
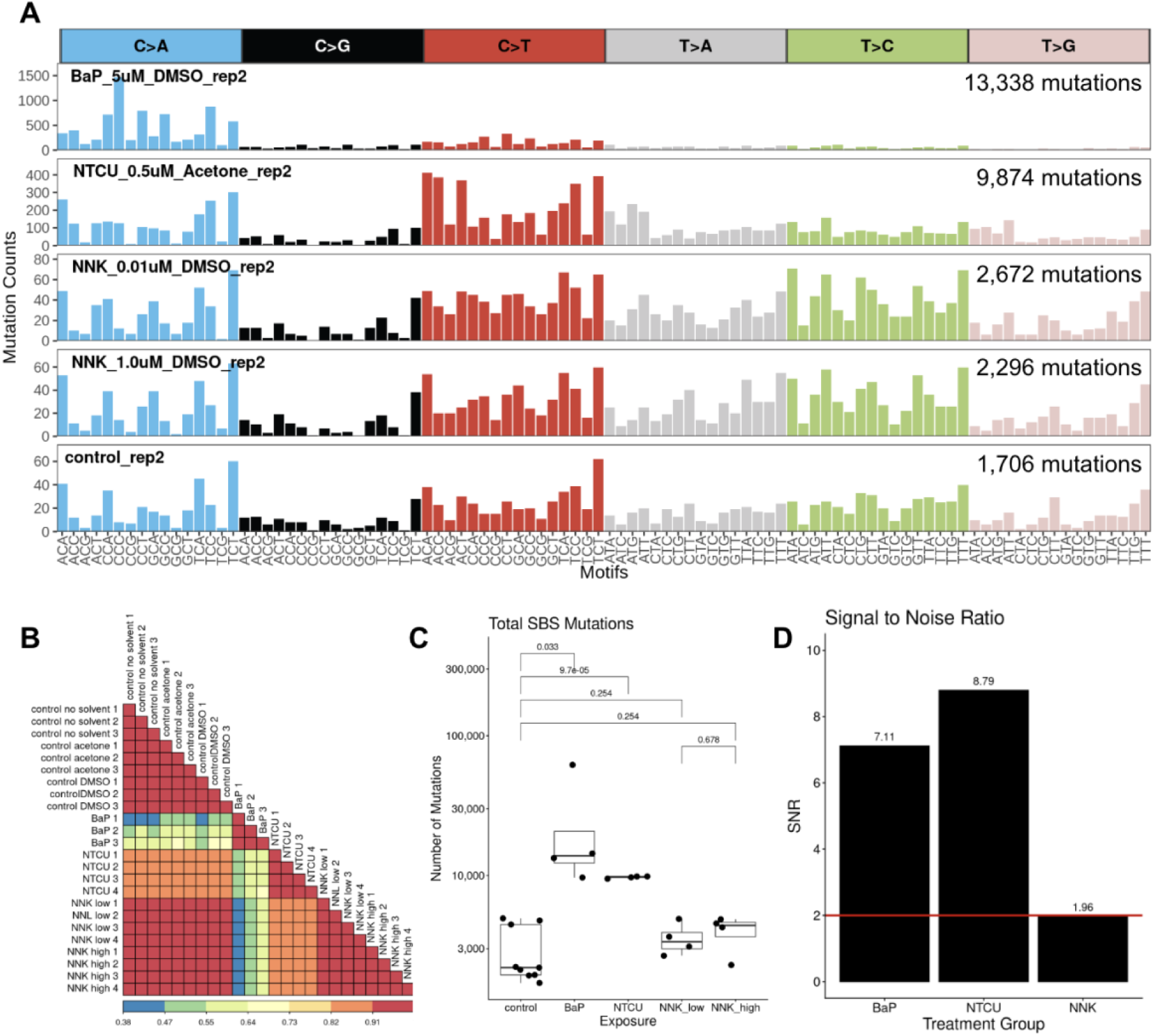
SBS patterns for BaP and NTCU treated samples are distinct from controls. **(A)** One representative SBS mutational pattern is shown for each of the four treatment groups and the controls. **(B)** The cosine correlation metric was used to assess the similarity of SBS patterns within and between conditions. High correlation existed between replicates within each group (cosine > 0.9). The pattern of SBS in NNK samples was highly correlated to controls (cosine > 0.9) while BaP and NTCU were less strongly correlated (cosine < 0.85). **(C)** The total number of SBS mutations is shown by condition. T-tests on log transformed data was used to compare each treatment group to the controls. P-values were corrected for multiple hypothesis testing using with Benjamini-Hochberg False Discovery Rate. The BaP and NTCU groups had significantly more SBSs compared to the control group. **(D)** The signal to noise ratio (SNR) is shown for each treatment group of mutations compared to the controls. An SNR above 2 (red line) suggests substantial dissimilarity between the background pattern of mutation and the treatment pattern. The mutational profiles of BaP and NTCU had large SNRs (>7) while the SNR for NNK was lower (1.96).

The number of SNVs in treated samples were compared to controls to determine which treatments led to higher mutation rates (**Figure 2C**). The average mutational loads were 24,826 for BaP, 9,735 for NTCU, 3,790 for NNK, and 2,925 for the control group. Pairwise comparisons were performed on log-transformed mutation counts with Benjamini–Hochberg correction. Compared to controls, BaP treatment resulted in significantly higher mutation counts (2.78 log2 fold change increase, 95% CI: 0.98-4.58, adjusted p = 0.033). NTCU treatment was also associated with a significant increase in mutation counts relative to control (1.86 log2 fold change increase, 95% CI: 1.38-2.35, adjusted p = 0.000097). No differences beyond random chance were found between either NNK concentration and the control group. Furthermore, there was no difference between the samples treated with low and high concentrations of NNK. Signal to noise ratio (SNR) was used to determine which mutational profiles were distinct from the control, or background, pattern of mutation (**Figure 2D**). SNR depends on the mean Euclidean distance between the mutational profiles of control and treatment samples (signal), as well as the variability within the control group and within the treatment group (noise). A large SNR value indicates the mutational pattern is distinct from the background pattern, since the distance between the two groups is distinguishable from the noise within each group [6]. BaP and NTCU samples showed a substantial signal from the background pattern (SNR > 7) while the NNK samples exhibited less signal (SNR = 1.96). Overall, these results indicate that BaP and NTCU treated samples exhibited a higher number and distinct pattern of SBSs compared to controls.

### Identification of the mutational signature for NTCU

To determine the mutational signatures associated with the carcinogens, musicatk was used to extract three *de novo* signatures from all 25 treatment and control samples (**Figure 3A**). Signature 1 was dominated by C>A mutations, specifically at the C[C>A]C context and resembles the pattern of mutations observed in the BaP treated samples. Signature 2 had mutations from all six SBS types and showed similarity to the pattern of mutations observed in the control samples and the NNK treated samples. Signature 3 also has mutations from all six substitution types present but with higher probability of C>T mutations and resembled the pattern of mutations observed in the NTCU treated samples.

**Figure 3.**
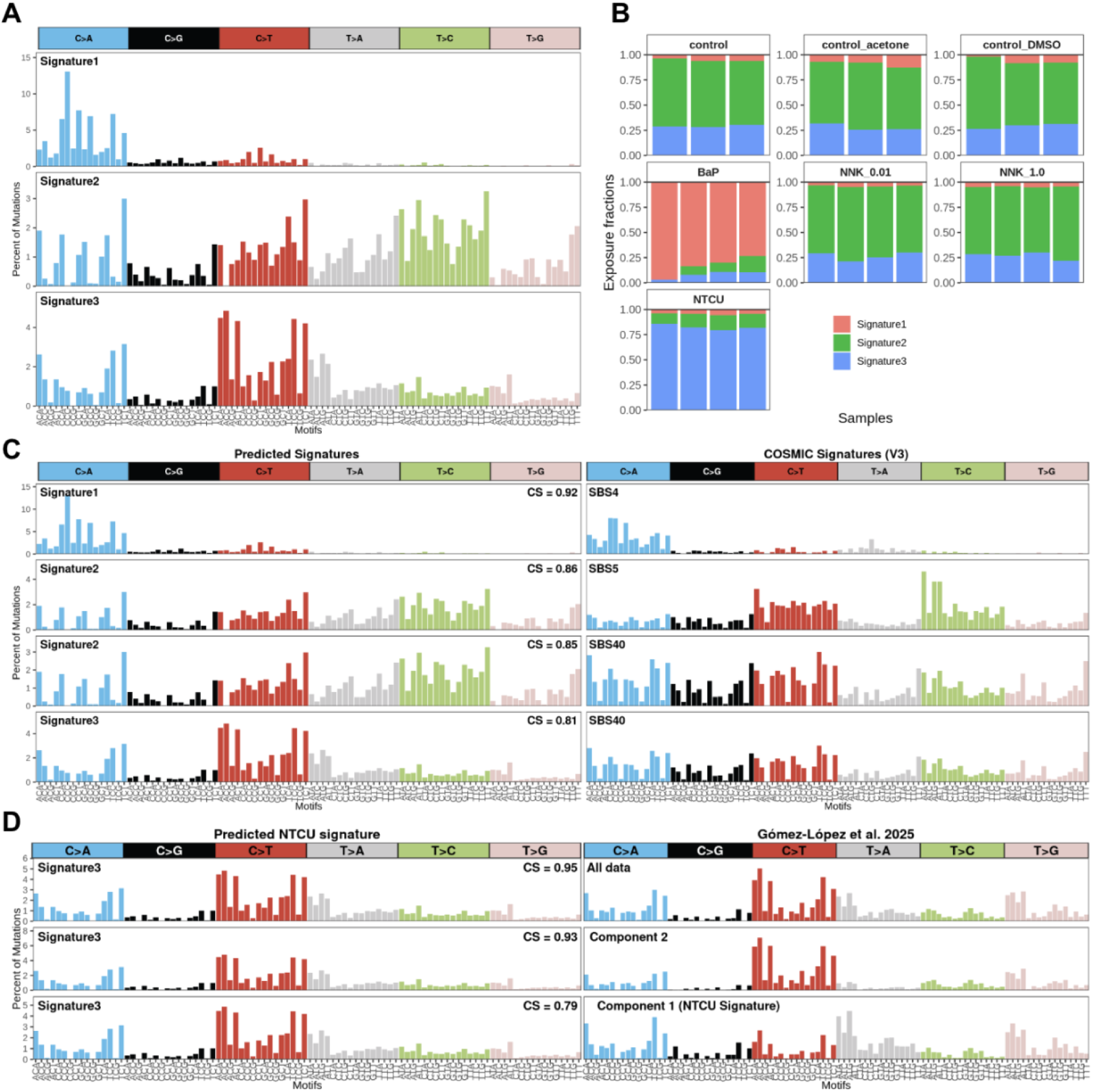
Deconvolution of SBS signatures from treated AALEs. **(A)** Three de novo signatures predicted by NMF using the musicatk package. **(B)** The proportion of mutations attributed to each of the three predicted signatures for each sample are shown and organized by condition. Mutations in BaP samples are nearly all attributed to Signature 1, which is not highly active in other conditions. Mutations in NTCU samples are nearly all attributed to Signature 3, which was not highly active in other conditions. **(C)** Comparison of predicted signatures to COSMIC v3 database. Signature 1 is the only signature with high cosine similarity to an existing COSMIC signature (cosine > 0.9). Signature 2 was modestly correlated to SBS5 and SBS40, which are “clock-like” signatures, suggesting it likely captures background mutational processes. Signature 3 had the lowest correlation to any COSMIC signature. **(D)** Comparison of the predicted NTCU signature to data from Gómez-López et al [24]. The’all data’ signature represents the aggregate mutational profile extracted as a single component. Components 1 and 2 result from deconvolving the full dataset into two signatures, with the authors designating Component 1 as the primary NTCU signature.

Quantifying the mutational contribution of each signature per sample revealed distinct exposure profiles for the BaP and NTCU treatment groups (**Figure 3B**). Mutations in BaP-exposed samples were overwhelmingly driven by Signature 1 (average 22,431 mutations; 90.4%), with minimal contributions from Signatures 2 (4.1%) and 3 (5.5%). Similarly, NTCU-treated samples exhibited a highly specific profile dominated by Signature 3, which accounted for an average of 8,016 mutations (82.3%), compared to just 4.7% and 13.0% for Signatures 1 and 2, respectively. In contrast, the signature compositions for both low-and high-dose NNK treatments were nearly indistinguishable from control samples. Both NNK groups and controls were primarily composed of Signature 2 (∼65–70%) and Signature 3 (∼26–29%), with Signature 1 contributing less than 6% of the mutational burden across all three groups.

To determine if the predicted signatures have similarity to documented signatures, the predicted signatures were compared to the COSMIC database of signatures (v3.0) using cosine correlation (**Figure 3C**). Predicted Signature 1, which drove the BaP treatment profile, strongly resembled COSMIC signature SBS4 (cosine = 0.92), a well-established marker of BaP and tobacco smoke exposure [1,6]. Conversely, Predicted Signature 2 (active in control and NNK samples) and Predicted Signature 3 (specific to NTCU treatment) lacked definitive matches to known etiological COSMIC signatures. While Signature 2 showed weak correlation to “clock-like” background signatures (e.g., SBS5; maximum cosine = 0.86), Signature 3 remained highly distinct (maximum cosine = 0.81).

To further validate this novel NTCU signature, we compared it to findings from a recent 2025 study by Gómez-López et al., which characterized an NTCU mutational signature using whole-genome sequencing of topically treated mouse epithelial cell [24]. Our *in vitro* human NTCU signature demonstrated a remarkably high correlation with the overall, aggregate mutational profile of their mouse data (cosine = 0.95; **Figure 3D**). Notably, while Gómez-López *et al*. were able to deconvolve their dataset into two distinct sub-signatures (reporting Component 1 as their primary NTCU signature), our single *in vitro* signature effectively captures the comprehensive mutational landscape of their *in vivo* model. The extraction of a single, encompassing signature in our analysis instead of two distinct components is likely due to the smaller number of NTCU-treated samples in our *in vitro* dataset, which naturally limits the statistical power required for NMF algorithms to further deconvolve these concurrent mutational patterns.

To validate our findings with an orthogonal approach, we applied BayesPowerNMF, a robust Bayesian modeling workflow that is highly sensitive and resistant to model misspecification (**Figure S4**) [25]. Reassuringly, this method consistently recapitulated the three primary signatures that closely resembled the mutational profiles of the exposure groups. However, BayesPowerNMF additionally extracted a fourth, low-abundance signature undetected by standard NMF. This minor signature did not reflect the overall mutational profiles of any specific treatment group, accounting for an average of only 73 mutations per sample. It was found in the highest quantities within the control samples (comprising 0–13% of their mutations) and showed similarity to COSMIC signature SBS18, suggesting it likely captures a low-level background mutational process, such as *in vitro* oxidative damage.

### NTCU and NNK lack specific DBS and INDEL signatures

To investigate double base substitution patterns, we utilized musicatk to extract three *de novo* signatures across all 25 samples (**Figure S5A**). Signature 1 exhibited high specificity for CC>AA and TG>AT mutations and was highly active exclusively in BaP-treated samples (**Figure S5B**). In these samples, Signature 1 accounted for an average of 290 mutations, representing 90% of the DBS burden, with minimal activity observed in any other group. Validating this finding, Signature 1 was the only profile demonstrating strong similarity to a known COSMIC signature, specifically matching the BaP-associated DBS2 (cosine = 0.91; **Figure S5C**). Conversely, Signatures 2 and 3 displayed flatter, non-specific mutational profiles. While NTCU samples were dominated by Signature 2 (averaging 69 mutations; 86% of the burden), this profile was not specific to NTCU, as it was also highly active across several control and NNK replicates.

Furthermore, Signature 3 activity was sporadic among control and NNK samples, reflecting highly inconsistent exposure compositions across these groups.

Similarly, we evaluated insertion and deletion mutational patterns by extracting three *de novo* signatures across all 25 samples (**Figure S6A**). Signature 2, characterized predominantly by single base pair deletions of C, was highly specific to the BaP cohort (**Figure S6B**). This signature accounted for roughly half of the mutations in three BaP replicates and nearly all mutations in the fourth. Validating this profile, Signature 2 strongly resembled COSMIC signature ID3 (cosine = 0.93), a pattern previously associated with BaP exposure (**Figure S6C**). In contrast, Signature 1 showed specific peaks at 5+ base pairs deletion (1 repeat), 5+ base pairs insertion (zero repeats), and in the microhomology deletion category, while Signature 3 displayed specific peaks at single base pair deletion of T (6+ homopolymer length), single base pair insertion of T (5+ homopolymer length), and 5+ base pairs insertion (zero repeats). Signatures 1 and 3 lacked treatment specificity as the signature contributions were comparable across the control, NNK, and NTCU groups. Overall, these results show that while BaP exhibits specific mutational activity, NTCU and NNK do not induce consistent or specific mutational signatures in the DBS or INDEL contexts.

## DISCUSSION

In this study, we characterized the previously undefined mutational signatures of two potent carcinogens widely utilized to model tobacco smoke-associated diseases. To achieve this, we developed a novel, physiologically relevant system combining human airway epithelial cells with whole-genome sequencing to assess mutational profiles induced by lung cancer-associated genotoxins. Previous whole-genome sequencing studies of normal bronchial epithelium have demonstrated a substantial tobacco smoke-induced mutation burden, supporting the use of these cells as a robust model for investigating lung carcinogens [26,27]. Importantly, our analysis of BaP-exposed samples yielded results highly consistent with existing literature, fully validating the accuracy of this platform. Compared to non-human *in vivo* models such as *C. elegans* or mice, our system offers direct human translational relevance. Furthermore, *in vitro* exposure provides precise experimental control over dosage and duration, minimizing confounding factors and allowing discovered signatures to be causally linked to a single exposure with high confidence.

Using this platform, we identified a novel mutational pattern definitively associated with NTCU exposure. NTCU-treated cells exhibited a significantly elevated mutational burden characterized by a distinct profile of acquired single base substitutions (SBSs). Specifically, our deconvolution extracted an SBS signature that was highly active in, and specific to, NTCU-exposed samples. Because this signature lacked strong correlation with any established profile in the COSMIC catalogue, it represents a previously uncharacterized pattern of mutagenesis. Conversely, NTCU exposure did not induce distinct mutational patterns in the double base substitution or insertion/deletion contexts, as the signatures extracted in those schemas were driven by background activity present across both treated and control samples.

Analysis of NNK-exposed samples yielded no distinct mutational patterns above background, maintaining a similar mutational burden and profile to control samples across all tested concentrations and mutational schemas (SBS, DBS, INDEL). Rather than suggesting NNK does not induce these mutation types, this negative result likely highlights a limitation of the in vitro metabolic activation system. NNK is a pro-carcinogen requiring extensive alpha-hydroxylation by specific enzymes, predominantly CYP2A13 and CYP2A6, to form mutagenic DNA adducts [28,29]. While our system effectively bioactivated BaP via CYP1A1/1B1, the supplemented liver microsomes and AALE cells may not have provided adequate CYP2A activity to metabolize NNK into its DNA-reactive intermediates, underscoring the need for tailored metabolic consideration when utilizing *in vitro* models for specific nitrosamines.

Beyond our primary findings, the application of the BayesPowerNMF workflow highlighted the importance of orthogonal computational validation in *in vitro* toxicogenomics. While standard NMF effectively captured the dominant carcinogen-induced patterns, the heightened sensitivity of the Bayesian approach revealed a fourth, low-abundance mutational signature predominantly active in the control samples. This minor background profile exhibits strong concordance with a control-derived signature previously reported in similar *in vitro* models by Kucab et al. [6]. The independent identification of this background noise across different studies strongly supports its biological authenticity as a true cell-culture artifact, rather than a mathematical anomaly. Furthermore, its detection underscores the utility of robust Bayesian methods in isolating very low-activity mutational processes that may fall below the detection thresholds of standard bootstrap-based NMF methods.

Mutational signatures provide valuable insight into cancer development and etiology; however, the mutational patterns of many known carcinogens remain uncharacterized. To address this, we developed a novel *in vitro* system leveraging whole-genome sequencing of human airway epithelial cells. After validating the accuracy of this platform by successfully recapturing the known SBS4 signature associated with BaP, we defined the previously unknown SBS signature of NTCU, a potent inducer of lung squamous cell carcinoma. While NNK did not yield a distinct profile—potentially reflecting a lack of specific *in vitro* metabolic requirements—our positive findings demonstrate the robust utility of this model. With broader application, this controlled, human-relevant system can be utilized to definitively establish causal links between environmental exposures and cancer and elucidate the etiology of mutational patterns in cancer genomes.

## DATA AVAILABILITY STATEMENT

Mutation counts and analysis code is available on GitHub at https://github.com/campbio-manuscripts/NTCU_Mutsigs.

## ACKNOWLEDGMENTS

We thank Dr. Sam M. Janes, Dr. Moritz J. Przybilla, and Dr. Sandra Gómez-López for sharing data and details regarding their NTCU associated signature.

## SUPPLEMENTARY FIGURES

**Supplementary Figure 1.**
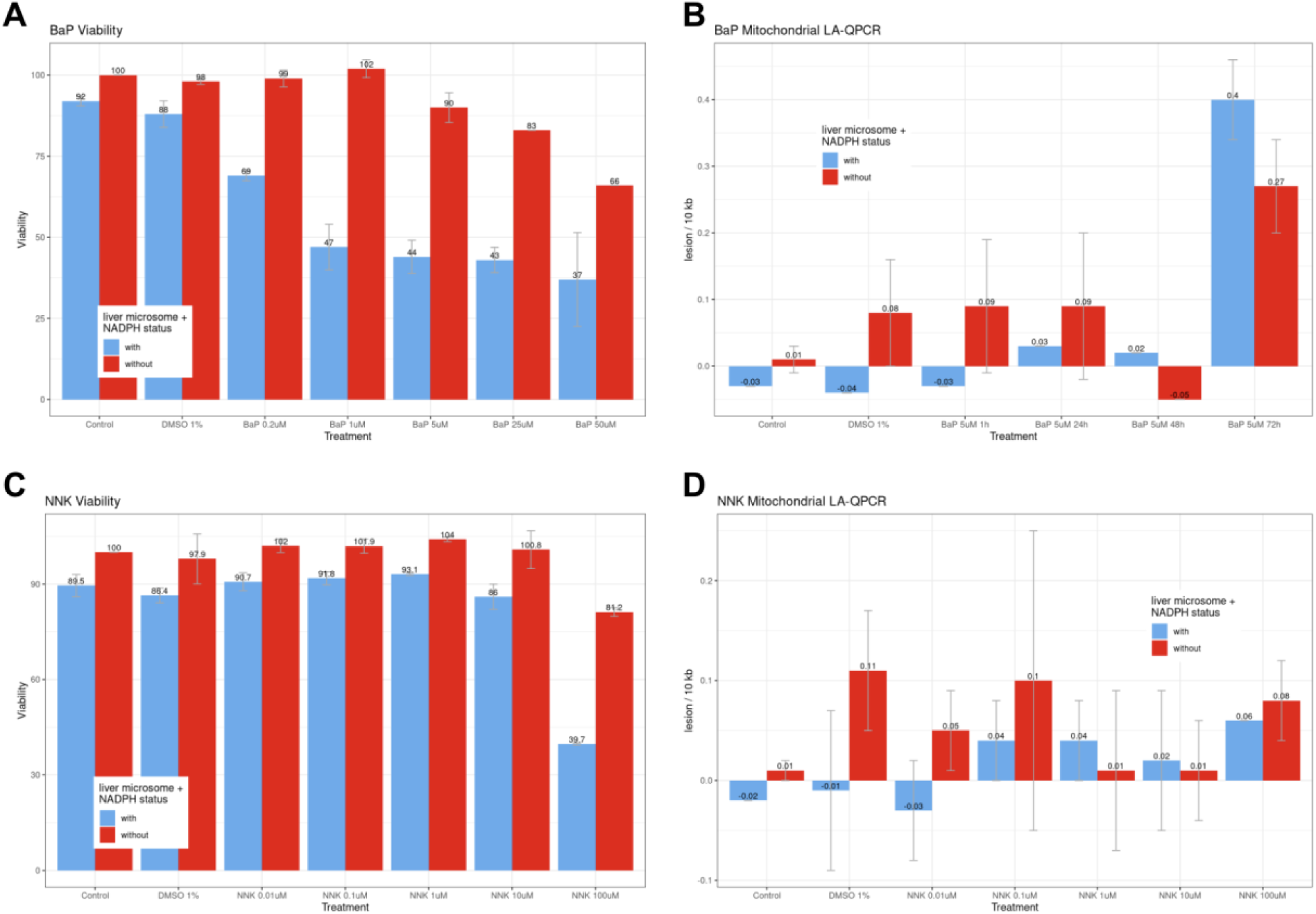
Cytotoxicity and DNA damage response analysis for BaP and NNK. **(A)** Cytotoxicity analysis results to determine concentration of BaP that results in 40-60% viability for seven days. Concentration of 5 μM was selected. **(B)** LA-QPCR analysis to determine DNA damage response. BaP with liver microsome causes more mitochondrial DNA damage than BaP alone. **(C)** Cytotoxicity analysis results to determine concentration of NNK that results in 40-60% viability for seven days. There was limited change in viability after NNK treatment, so two concentrations were selected: 0.01 μM and 1.0 μM. **(D)** LA-QPCR analysis to determine DNA damage response. NNK causes limited mitochondrial DNA damage with and without liver microsome, so NNK alone treatment was selected.

**Supplementary Figure 2.**
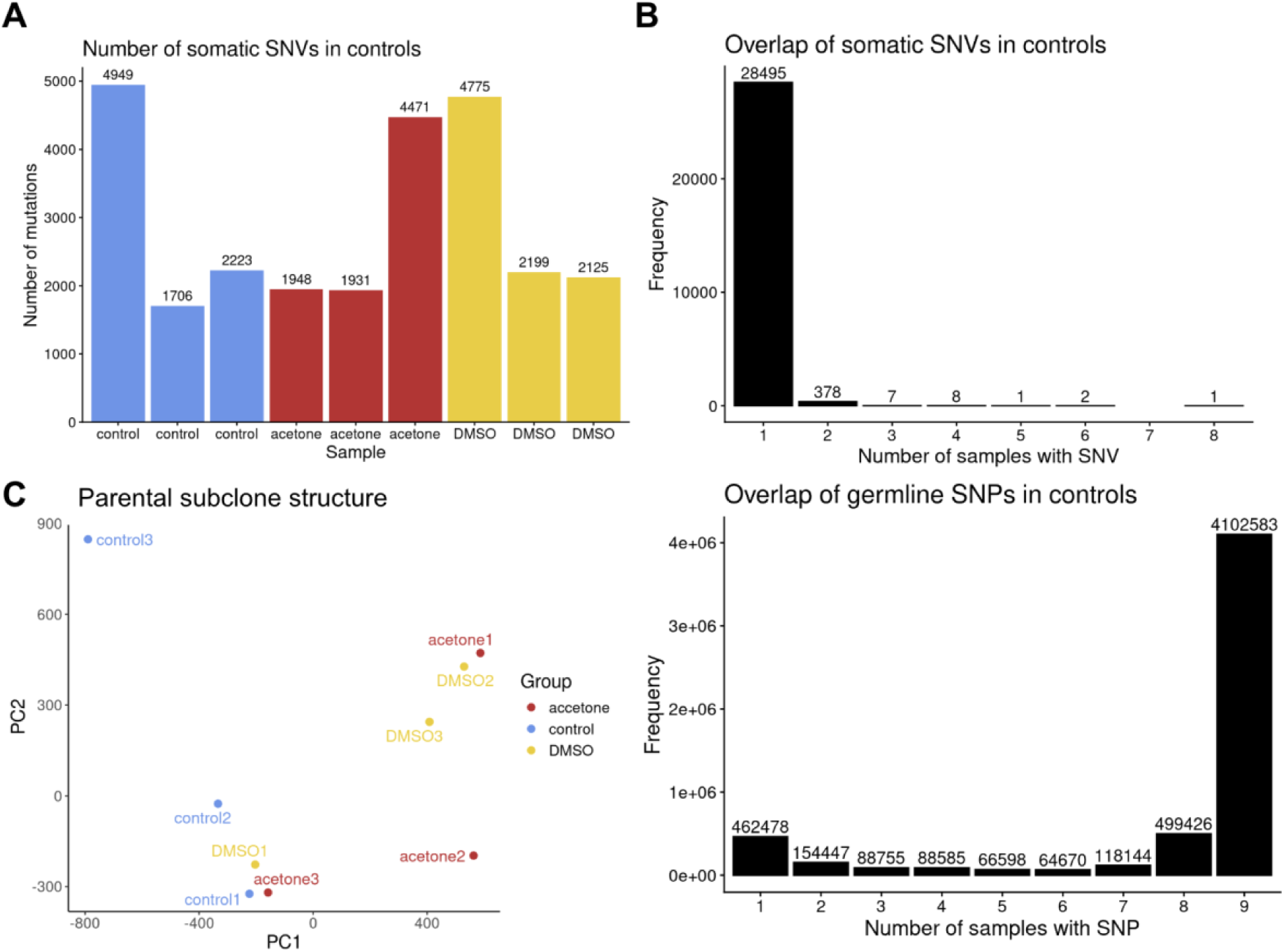
Analysis of subclonal structure in control samples. **(A)** Total number of SBS mutations in each control sample. Notably, one replicate per condition (control1, acetone3, and DMSO1) exhibits a markedly higher mutational burden. **(B)** Overlap of SNVs identified via the somatic mutation pipeline. As expected, nearly all somatic SNVs are unique to a single sample. **(C)** Principal component analysis (PCA) resolving the subclonal architecture of the control samples. To isolate subclonal structure, variants universally present in all nine samples (shared germline/parental clonal) and variants unique to a single sample (private somatic) were filtered out. PCA based on the remaining shared variants reveals three distinct clusters representing distinct parental subclones. Subclone 1 comprises acetone1, acetone2, DMSO2, and DMSO3; Subclone 2 comprises control2 and control3; and Subclone 3 comprises control1, acetone3, and DMSO1. **(D)** Overlap of germline SNPs across control samples. The vast majority of SNPs are present in all nine replicates, confirming a shared parental origin.

**Supplementary Figure 3.**
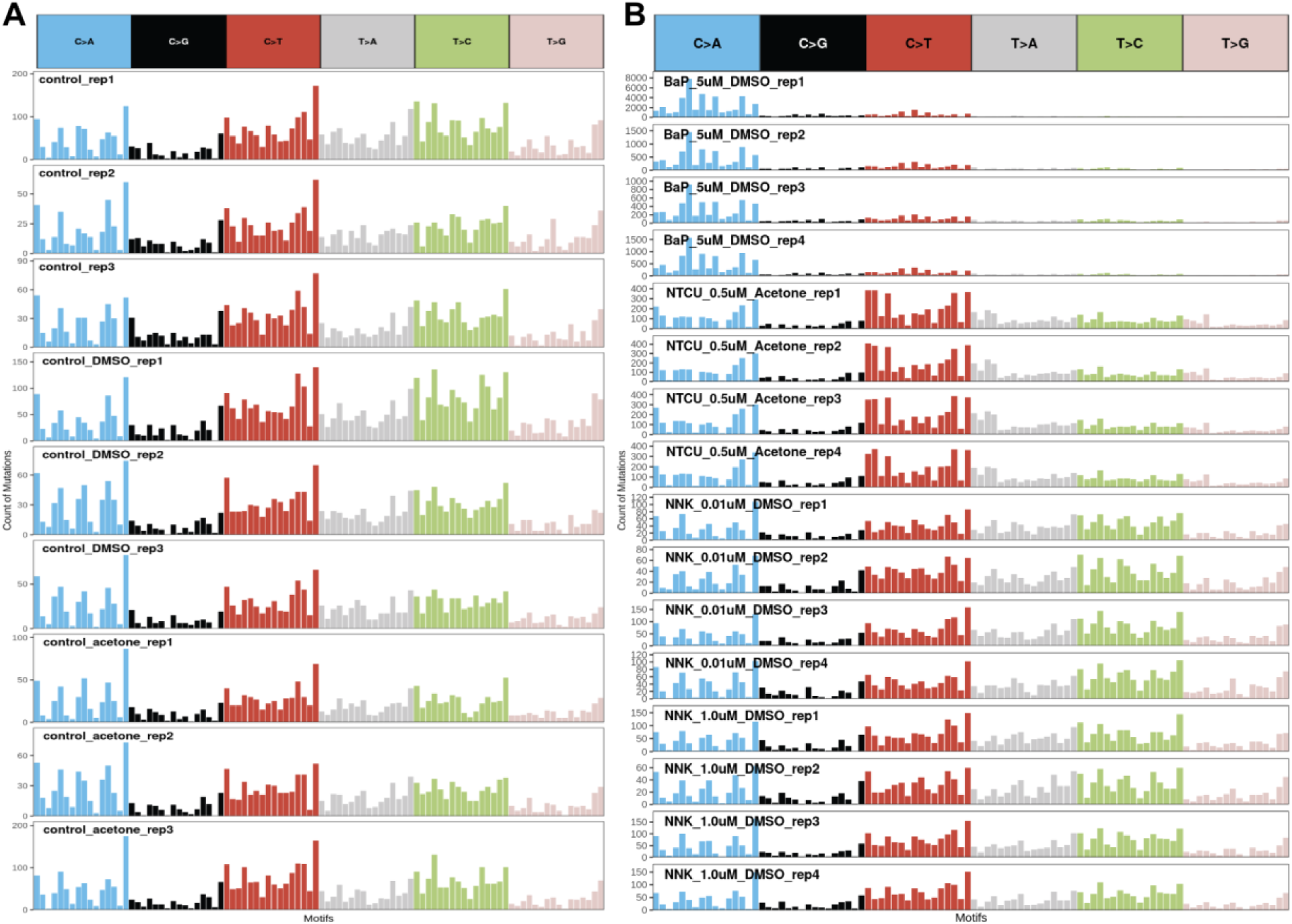
SBS mutation patterns of all samples. **(A)** Frequency of SBS mutation motifs in all control samples. **(B)** Frequency of SBS mutation motifs in all treatment samples.

**Supplementary Figure 4.**
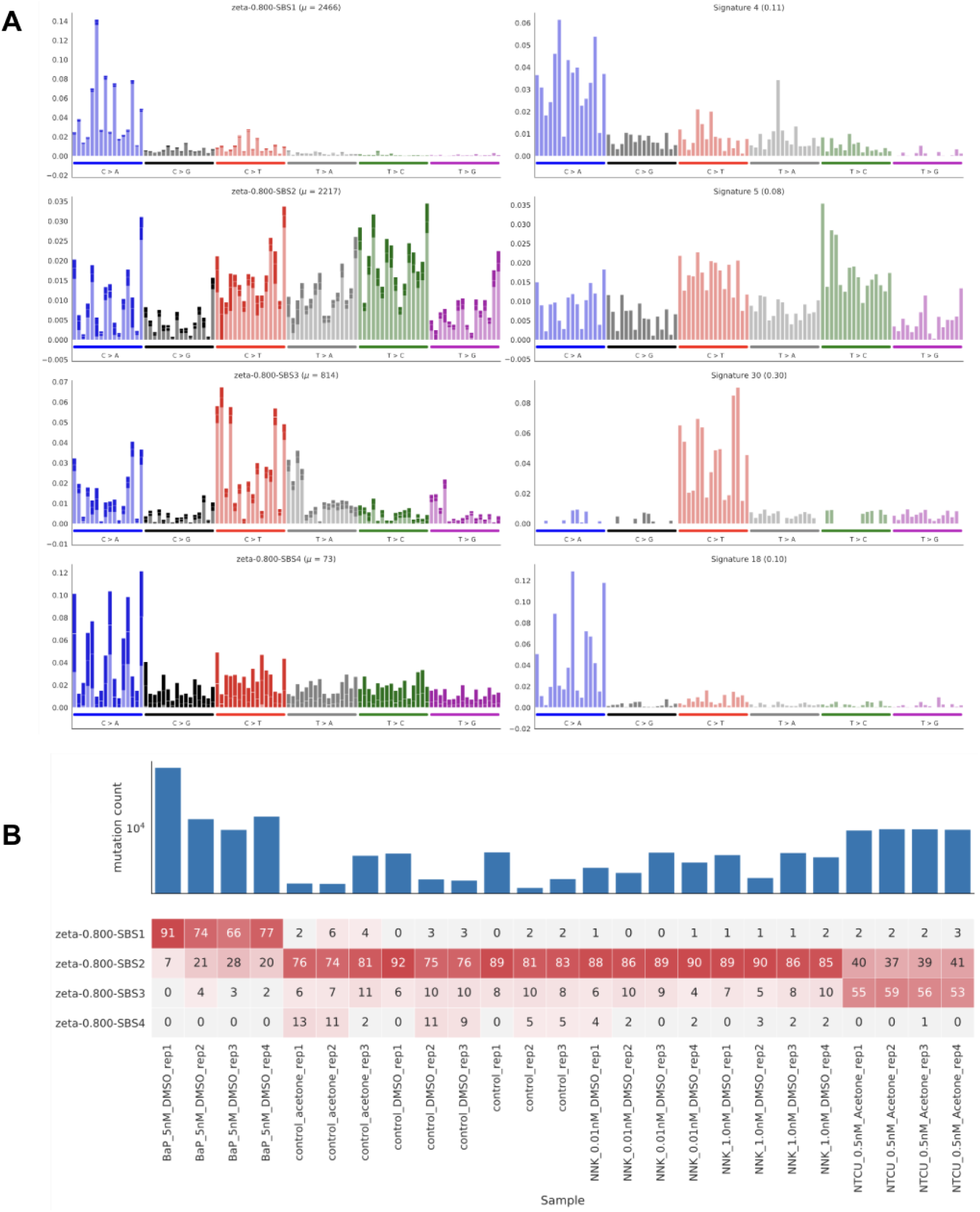
SBS signatures discovered with BayesPowerNMF. **(A)** Four signatures predicted by BayesPowerNMF (left) with their most similar COSMIC signature (right). A fourth signature not found by standard NMF workflows was present at lower levels. **(B)** Percent of mutations attributed to each of the four predicted signatures for each sample. Activity levels for the first three signatures are consistent with those from the NMF method. The new fourth signature accounted for 0-13% of mutations in the control samples and was highly correlated with COSMIC SBS18.

**Supplementary Figure 5.**
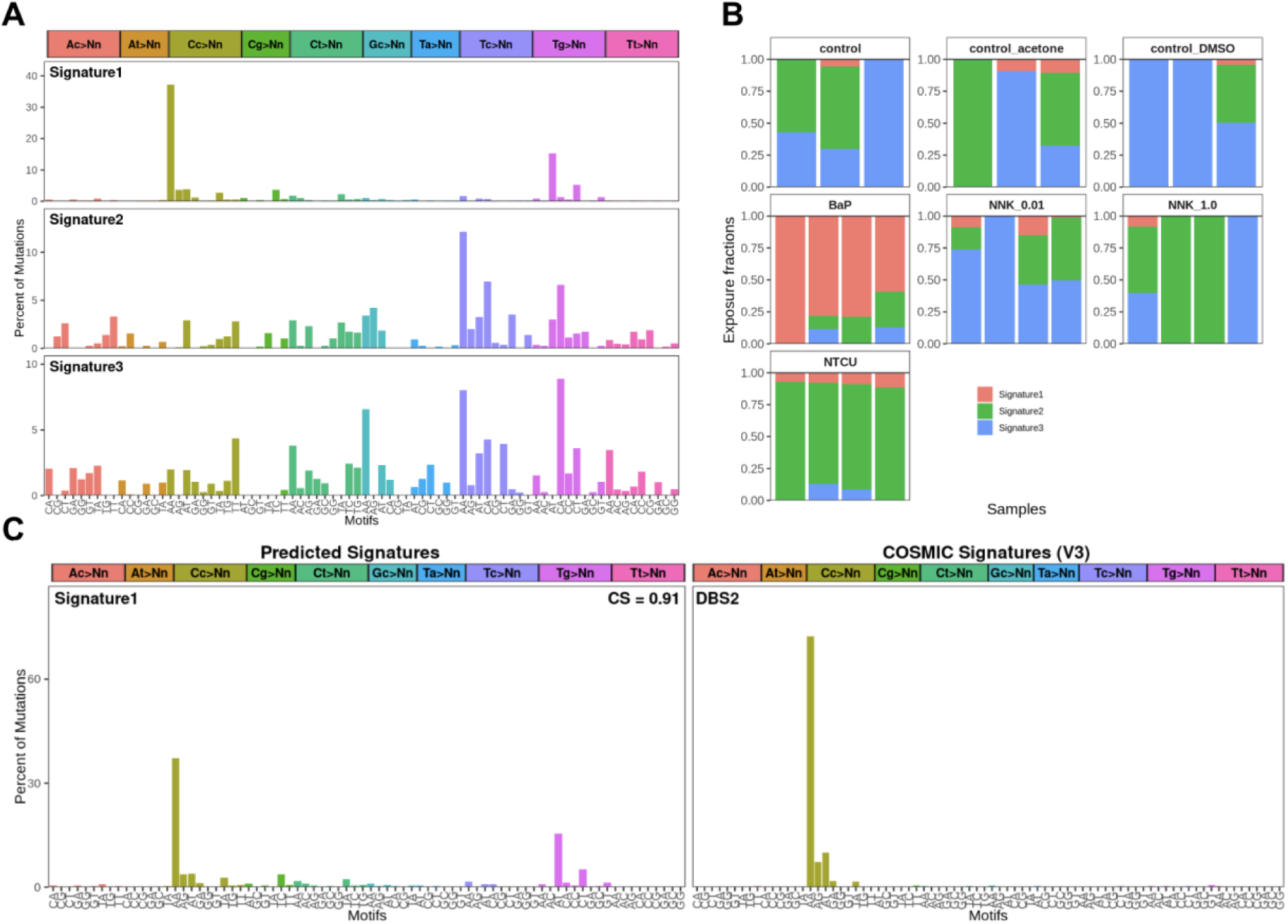
DBS signature analysis. **(A)** Three DBS signatures were predicted by NMF in the musicatk package. **(B)** Proportion of mutations attributed to each of the three predicted signatures for each sample, organized by group. Mutations in BaP samples are nearly all attributed to Signature 1, which is not highly active in any other sample type. Mutations in NTCU samples are nearly all attributed to Signature 2, but this signature is also highly active in other sample types. **(C)** Comparison of predicted signatures to COSMIC v3 database. Signature 1 is the only signature with relatively high cosine similarity to an existing signature (cosine > 0.9).

**Supplementary Figure 6.**
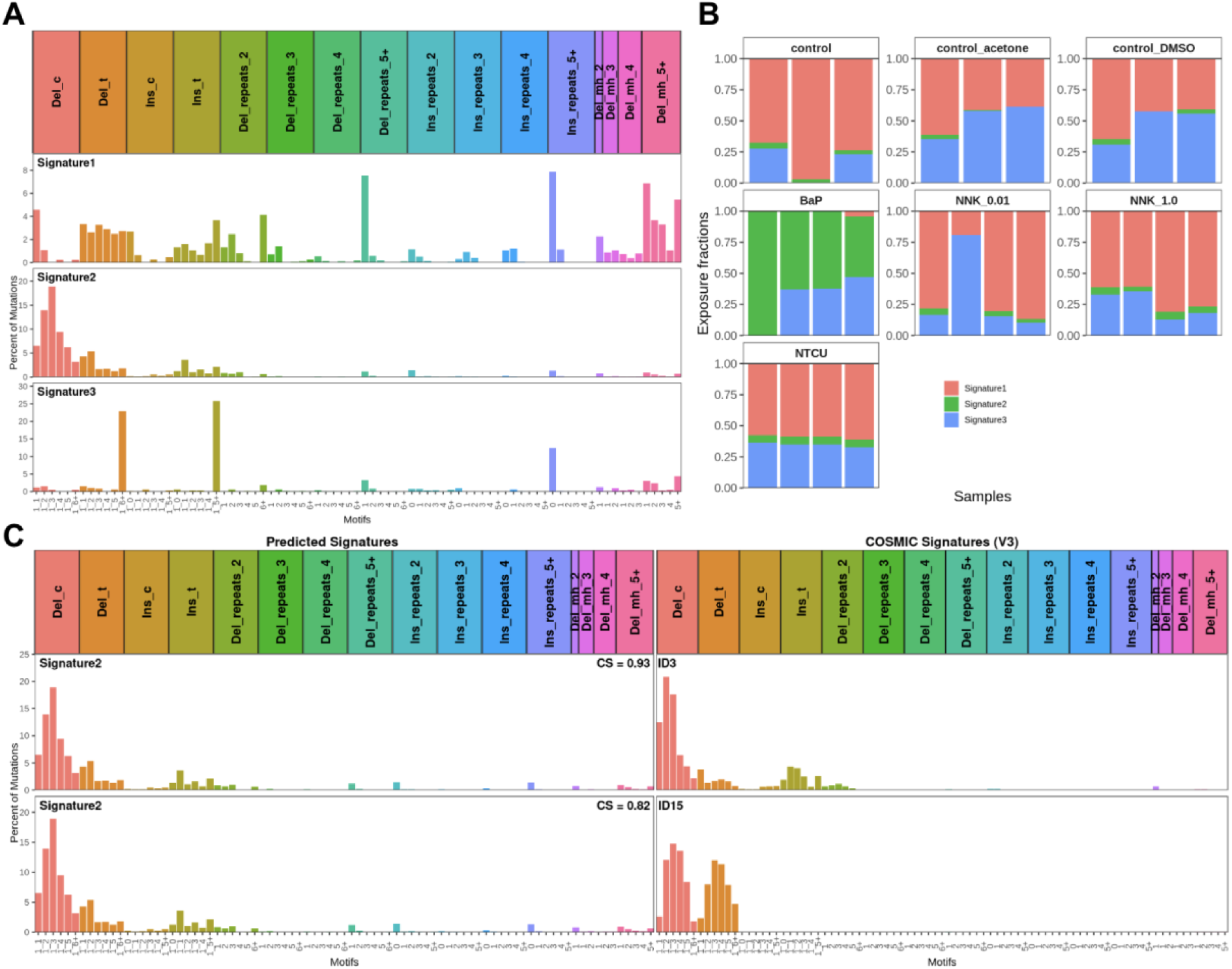
INDEL signature analysis. **(A)** Three INDEL signatures predicted by NMF in the musicatk package. **(B)** Proportion of mutations attributed to each of the three predicted signatures for each sample, organized by group. Signature 2 is highly active only in BaP samples. **(C)** Comparison of predicted signatures to COSMIC v3 database. Signature 2 is the only signature with relatively high cosine similarity to an existing signature (cosine > 0.9).

## REFERENCES

1. Alexandrov LB, Nik-Zainal S, Wedge DC et al. Signatures of mutational processes in human cancer. Nature 2013;500:415–21.

2. Lawrence MS, Stojanov P, Polak P et al. Mutational heterogeneity in cancer and the search for new cancer-associated genes. Nature 2013;499:214–8.

3. Alexandrov LB, Kim J, Haradhvala NJ et al. The repertoire of mutational signatures in human cancer. Nature 2020;578:94–101.

4. Nik-Zainal S, Kucab JE, Morganella S et al. The genome as a record of environmental exposure. Mutagenesis 2015;30:763–70.

5. Tate JG, Bamford S, Jubb HC et al. COSMIC: the Catalogue Of Somatic Mutations In Cancer. Nucleic Acids Res 2019;47:D941–7.

6. Kucab JE, Zou X, Morganella S et al. A Compendium of Mutational Signatures of Environmental Agents. Cell 2019;177:821–836.e16.

7. Huskova H, Ardin M, Weninger A et al. Modeling cancer driver events in vitro using barrier bypass-clonal expansion assays and massively parallel sequencing. Oncogene 2017;36:6041–8.

8. Schraven B, Roux M, Hutmacher B et al. Triggering of the alternative pathway of human T cell activation involves members of the T 200 family of glycoproteins. Eur J Immunol 1989;19:397–403.

9. Mazzilli SA, Hershberger PA, Reid ME et al. Vitamin D Repletion Reduces the Progression of Premalignant Squamous Lesions in the NTCU Lung Squamous Cell Carcinoma Mouse Model. Cancer Prev Res (Phila) 2015;8:895–904.

10. Xiong D, Pan J, Yin Y et al. Novel mutational landscapes and expression signatures of lung squamous cell carcinoma. Oncotarget 2018;9:7424–41.

11. Wang Y, Zhang Z, Garbow JR et al. Chemoprevention of lung squamous cell carcinoma in mice by a mixture of Chinese herbs. Cancer Prev Res (Phila) 2009;2:634–40.

12. Belinsky SA, Devereux TR, Foley JF et al. Role of the alveolar type II cell in the development and progression of pulmonary tumors induced by 4-(methylnitrosamino)-1-(3-pyridyl)-1-butanone in the A/J mouse. Cancer Res 1992;52:3164–73.

13. Hecht SS, Morse MA, Amin S et al. Rapid single-dose model for lung tumor induction in A/J mice by 4-(methylnitrosamino)-1-(3-pyridyl)-1-butanone and the effect of diet. Carcinogenesis 1989;10:1901–4.

14. Hecht SS, Isaacs S, Trushin N. Lung tumor induction in A/J mice by the tobacco smoke carcinogens 4-(methylnitrosamino)-1-(3-pyridyl)-1-butanone and benzo[a]pyrene: a potentially useful model for evaluation of chemopreventive agents. Carcinogenesis 1994;15:2721–5.

15. Hecht SS. Tobacco smoke carcinogens and lung cancer. J Natl Cancer Inst 1999;91:1194–210.

16. Schuller HM, Tithof PK, Williams M et al. The tobacco-specific carcinogen 4-(methylnitrosamino)-1-(3-pyridyl)-1-butanone is a beta-adrenergic agonist and stimulates DNA synthesis in lung adenocarcinoma via beta-adrenergic receptor-mediated release of arachidonic acid. Cancer Res 1999;59:4510–5.

17. Meyer JN. QPCR: a tool for analysis of mitochondrial and nuclear DNA damage in ecotoxicology. Ecotoxicology 2010;19:804–11.

18. Broad Institute GR. Picard Toolkit. 2019.

19. Van Der Auwera GA, Carneiro MO, Hartl C et al. From FastQ Data to High-Confidence Variant Calls: The Genome Analysis Toolkit Best Practices Pipeline. CP in Bioinformatics 2013;43, DOI: 10.1002/0471250953.bi1110s43.

20. Benjamin D, Sato T, Cibulskis K et al. Calling Somatic SNVs and Indels with Mutect2. 2019, DOI: 10.1101/861054.

21. Chevalier A, Yang S, Khurshid Z et al. The Mutational Signature Comprehensive Analysis Toolkit (musicatk) for the Discovery, Prediction, and Exploration of Mutational Signatures. Cancer Res 2021;81:5813–7.

22. Poplin R, Ruano-Rubio V, DePristo MA et al. Scaling accurate genetic variant discovery to tens of thousands of samples. 2017, DOI: 10.1101/201178.

23. Islam SMA, Díaz-Gay M, Wu Y et al. Uncovering novel mutational signatures by de novo extraction with SigProfilerExtractor. Cell Genom 2022;2:None.

24. Gómez-López S, Alhendi ASN, Przybilla MJ et al. Aberrant basal cell clonal dynamics shape early lung carcinogenesis. Science 2025;388:eads9145.

25. Xue C, Miller JW, Carter SL et al. Robust discovery of mutational signatures using power posteriors. bioRxiv 2024:2024.10.23.619958.

26. Yoshida K, Gowers KHC, Lee-Six H et al. Tobacco smoking and somatic mutations in human bronchial epithelium. Nature 2020;578:266–72.

27. Spencer Chapman M, Mitchell E, Yoshida K et al. Prolonged persistence of mutagenic DNA lesions in somatic cells. Nature 2025;638:729–38.

28. Su T, Bao Z, Zhang QY et al. Human cytochrome P450 CYP2A13: predominant expression in the respiratory tract and its high efficiency metabolic activation of a tobacco-specific carcinogen, 4-(methylnitrosamino)-1-(3-pyridyl)-1-butanone. Cancer Res 2000;60:5074–9.

29. He X-Y, Shen J, Ding X et al. Identification of critical amino acid residues of human CYP2A13 for the metabolic activation of 4-(methylnitrosamino)-1-(3-pyridyl)-1-butanone, a tobacco-specific carcinogen. Drug Metab Dispos 2004;32:1516–21.

